# The combination of morphogenic regulators BABY BOOM and GRF-GIF improves maize transformation efficiency

**DOI:** 10.1101/2022.09.02.506370

**Authors:** Zongliang Chen, Juan M. Debernardi, Jorge Dubcovsky, Andrea Gallavotti

## Abstract

Transformation is an indispensable tool for plant genetics and functional genomic studies. Although stable transformation no longer represents a major technology bottleneck in maize, there is still need for easily accessible and efficient transformation methods in most academic labs. Here we present the GGB transformation system, a rapid and highly efficient transformation system optimized for the immature embryo transformation of two maize genetic backgrounds, including the inbred line B104. The combination of distinct morphogenetic factors, the maize BABY BOOM transcriptional regulator (ZmBBM/EREB53) and the wheat GRF4-GIF1 (GROWTH REGULATING FACTOR4 - GRF-INTERACTING FACTOR1) chimera, together with a modified QuickCorn protocol, regenerated transformed maize seedlings in approximately two months with an efficiency of 26 to 37%; notably, the efficiency was 7-fold higher than with using either component in isolation. Additionally, ectopic expression of both morphogenetic factors did not show obvious effects on B104 development, and in particular fertility was not affected, obviating the need to remove the morphogenetic regulators post *Agrobacterium* infections. The GGB transformation system is designed for CRISPR-Cas9 editing but can be adapted for other purposes and should be easy to implement in most academic labs with little transformation experience.

## INTRODUCTION

The promise of genome editing to rapidly advance crop improvement relies for most species on the ability to generate transformed plants. However, lengthy and inefficient methods for transformation and regeneration of recalcitrant species as well as the genotype-dependency of the transformation process hinder the widespread adoption of crop transformation technologies. For most species, simple transformation systems (e.g. floral dipping) are not a viable strategy, and tissue culture-based transformation systems involving callus-induction steps are the only possible avenue to regenerate transgenic plants. These however often carry unintended consequences (e.g. somaclonal variation) (Phillips *et al*. 1994; Neelakandan and wang 2012) and are not easy to implement.

In recent years, several technologies have been introduced to address these significant bottlenecks to crop transformation. These include the use of various morphogenic factors, genes that are involved in somatic embryogenesis or meristem development and that trigger reprogramming of a subset of somatic cells to eventually produce transformed plants (Gordon-kamm *et al*. 2019; Kausch *et al*. 2019; Maren *et al*. 2022). Inducing regeneration-competent or embryogenic cells is a critical step for plant transformation, and it has long been known that phytohormones, auxin and cytokinin, as well as wounding are the original triggering signals for plant regeneration. Key morphogenic regulators have been successfully manipulated to induce regeneration in crop plants. The most significant advancement was the introduction of the WUSCHEL2-BABY BOOM (WUS-BBM) system in maize and other crops by Corteva (Lowe *et al*. 2016; Lowe *et al*. 2018). The ectopic expression of the maize *BABY BOOM* gene (*ZmBBM/EREB53*) and *WUSCHEL2* (*ZmWUS2*) was first combined to increase transformation efficiency for maize transformation (Lowe *et al*. 2016). This system relies on the ectopic expression of two developmental regulators involved in meristem activity, the maize *ZmWUS2* and *ZmBBM/EREB53* genes. *ZmWUS2* is a maize co-ortholog of the Arabidopsis *WUS* gene (maize has another co-ortholog called *ZmWUS1*; (Nardmann and werr 2006). *WUS* is required for shoot apical meristem establishment during embryogenesis and for axillary meristem initiation post-embryogenesis (Mayer *et al*. 1998; Lenhard *et al*. 2002; Wang *et al*. 2017). *BBM* encodes an AP2/EREB transcription factor. Overexpression of *BBM* homologs in different plant species has been used to boost somatic embryogenesis in vegetative tissue (Boutilier *et al*. 2002; Deng *et al*. 2009; Heidmann *et al*. 2011). However, due to pleiotropic effects, removal of both morphogenic regulators from transgenic plants is required in this system (Lowe *et al*. 2018; Wang *et al*. 2020b). Alternative strategies have been pursued to obviate this issue (Lowe *et al*. 2018; Hoerster *et al*. 2020; Che *et al*. 2022). Recently, when combined with *Agrobacterium* strains carrying a helper plasmid (an improved ternary vector carrying additional virulence genes), another member of *WUSCHEL* family, *TaWOX5*, was found to dramatically increase transformation efficiency of the maize inbred lines B73 and A188, generating transgenic plants without obvious developmental defects (Wang *et al*. 2022b; Wang *et al*. 2022a).

The constitutive expression of the wheat *GRF4-GIF1* (*GROWTH REGULATING FACTOR4 - GRF-INTERACTING FACTOR1*) chimeric gene has also been shown to enhance transformation efficiency in wheat and other monocot species (Debernardi *et al*. 2020). Additionally, two individual maize GRFs (*ZmGRF5-LIKE1* and *ZmGRF5-LIKE2*) have been reported to boost transformation efficiency in the maize inbred A188 without pleiotropic effects in transgenic plants (Kong *et al*. 2020). As important developmental regulators, GRFs together with GIFs hold potential for improving transformation efficiency of different plant species (Debernardi *et al*. 2014; Debernardi *et al*. 2020). However, these transformation systems still require callus-inducing steps. Despite the recent breakthroughs using morphogenic factors, challenges still exist in developing fast, efficient and reliable transformation systems that can be quickly adopted by the public sector for basic and applied research, in particular for monocot species.

In this study, we combined the morphogenic regulators ZmBBM and TaGRF4-GIF1 in a single binary vector carrying Cas9 to explore the potential of this combination to enhance transformation efficiency in two maize genetic backgrounds, Hi-II and B104, commonly used by transformation facilities. While Hi-II is a mixed genetic background, which can complicate phenotypic analysis of transgenic plants, B104 is an inbred line related to B73, the first maize genome to be sequenced and assembled (Schnable *et al*. 2009; Raji *et al*. 2018; Aesaert *et al*. 2022; Kang *et al*. 2022). In combination with a modified QuickCorn protocol (Masters *et al*. 2020), we found that this system significantly enhanced the efficiency of transformation in both backgrounds, allowing the efficient generation of transgenic plants within a two-month time frame. We named this transformation system GGB, for GRF-GIF-BBM. We believe this rapid and highly efficient system can be adopted by most academic labs with little transformation experience. This study adds to a growing list of tools available for functional genomic studies in maize (Lowe *et al*. 2016; Mookkan *et al*. 2017; Lowe *et al*. 2018; Gordon-kamm *et al*. 2019; Hoerster *et al*. 2020; Masters *et al*. 2020; Kang *et al*. 2022).

## RESULTS

### The GGB system for maize transformation

Given that some morphogenic factors function in hierarchical order during embryogenesis and meristem formation (Wang *et al*. 2020a; WU *et al*. 2022), we reasoned that their combination may promote somatic embryogenesis in an additive manner, similar to the original BBM WUS2 system. We therefore tested whether the combination of the maize *BBM* gene (*ZmBBM/EREB53*) and the wheat GRF4-GIF1 chimera worked synergistically to enhance maize regeneration during *Agrobacterium*-mediated embryo transformation, as previously hypothesized (Debernardi *et al*. 2020). We first modified the binary plasmid pBUE411 (Xing *et al*. 2014) by building a construct containing *ZmBBM* expressed by the tissue specific promoter *pPLTP* (*PHOSPHOLIPID TRANSFERASE PROTEIN*) which promotes expression in embryos and leaves (Lowe *et al*. 2018; Jones *et al*. 2019), and the wheat GRF4-GIF1 chimera driven by the maize *UBIQUITIN1* promoter (Debernardi *et al*. 2020) (Figure 1). A CRISPR-Cas9 cassette and a *BsaI* cloning site for guide RNA (gRNA) cassettes are also included in the original pBUE411 vector for genome editing (pBUE411-GGB; Figure 1a). The original gRNA cassette in pBUE411 containing OsU3_promoter-SmR-gRNA_scaffold-OsU3_terminator was replaced by a short DNA fragment carrying double *BsaI* cut sites for single or multiple guide RNA cassette cloning, while the maize codon-optimized Cas9 gene driven by the *ZmUbi1* promoter was retained from the original vector.

**Figure 1.**
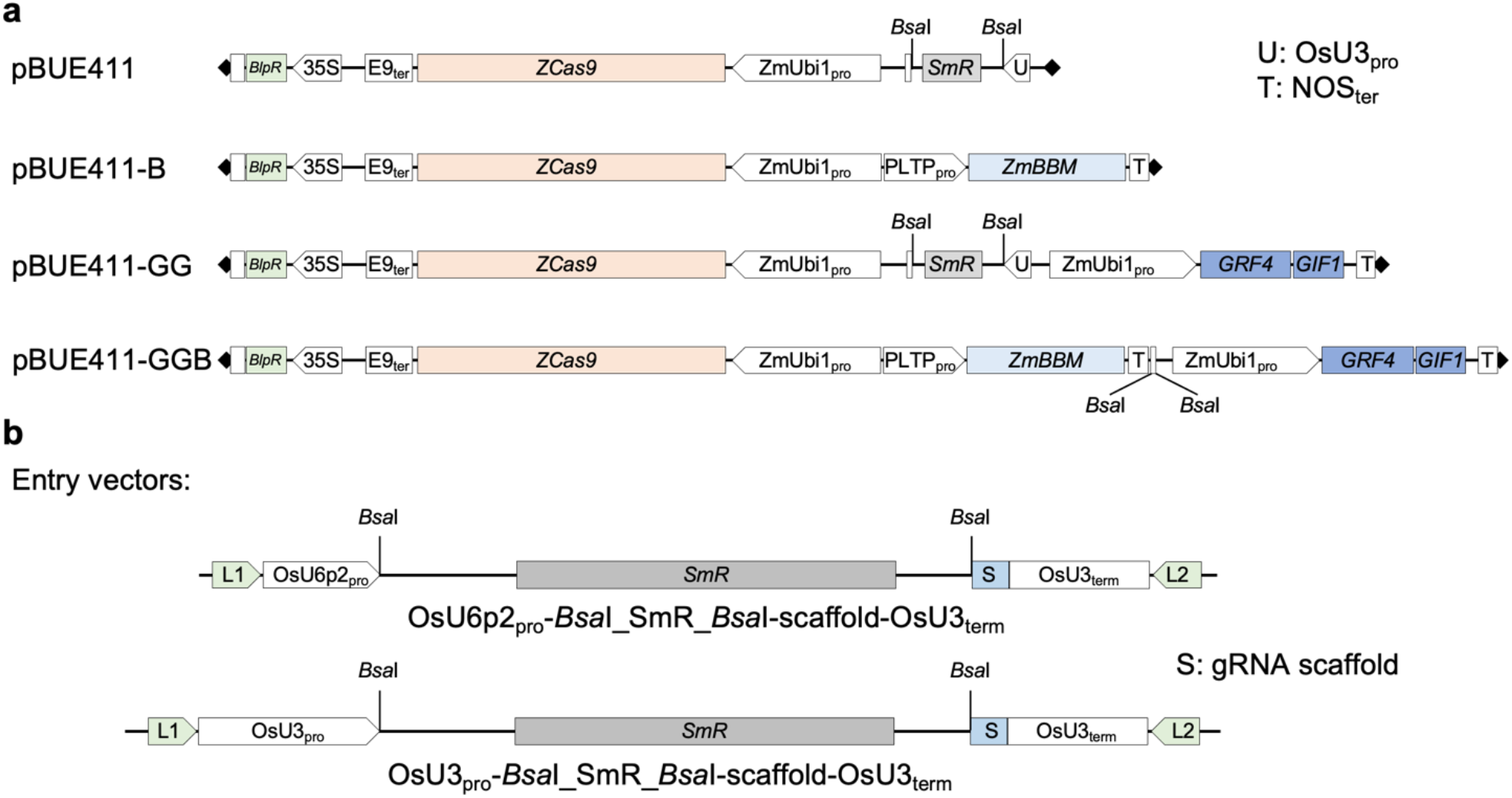
Construct maps. (**a**) Diagrams of four binary vectors used for stable maize transformation. (**b**) Diagrams of two entry vectors to generate individual gRNA cassettes.

Control transformations were carried out using the same vector backbone (pBUE411) carrying only the *BBM* (pBUE411-B) gene or the *GRF-GIF* chimera (pBUE411-GG) (Figure 1a). For the GGB system, we adopted the QuickCorn protocol developed by Corteva (Masters *et al*. 2020), reducing the time to obtain transformed plantlets to approximately 2 months (Figure 2e), compared to the 4-5 months usually required in standard transformation protocols (Frame *et al*. 2002), a timeframe comparable to the BBM WUS2 system (Lowe *et al*. 2018). The combination of both morphogenic factors and the utilization of the QuickCorn protocol yielded rapid and efficient somatic embryogenesis on immature embryos (Figures 2 and 3).

**Figure 2.**
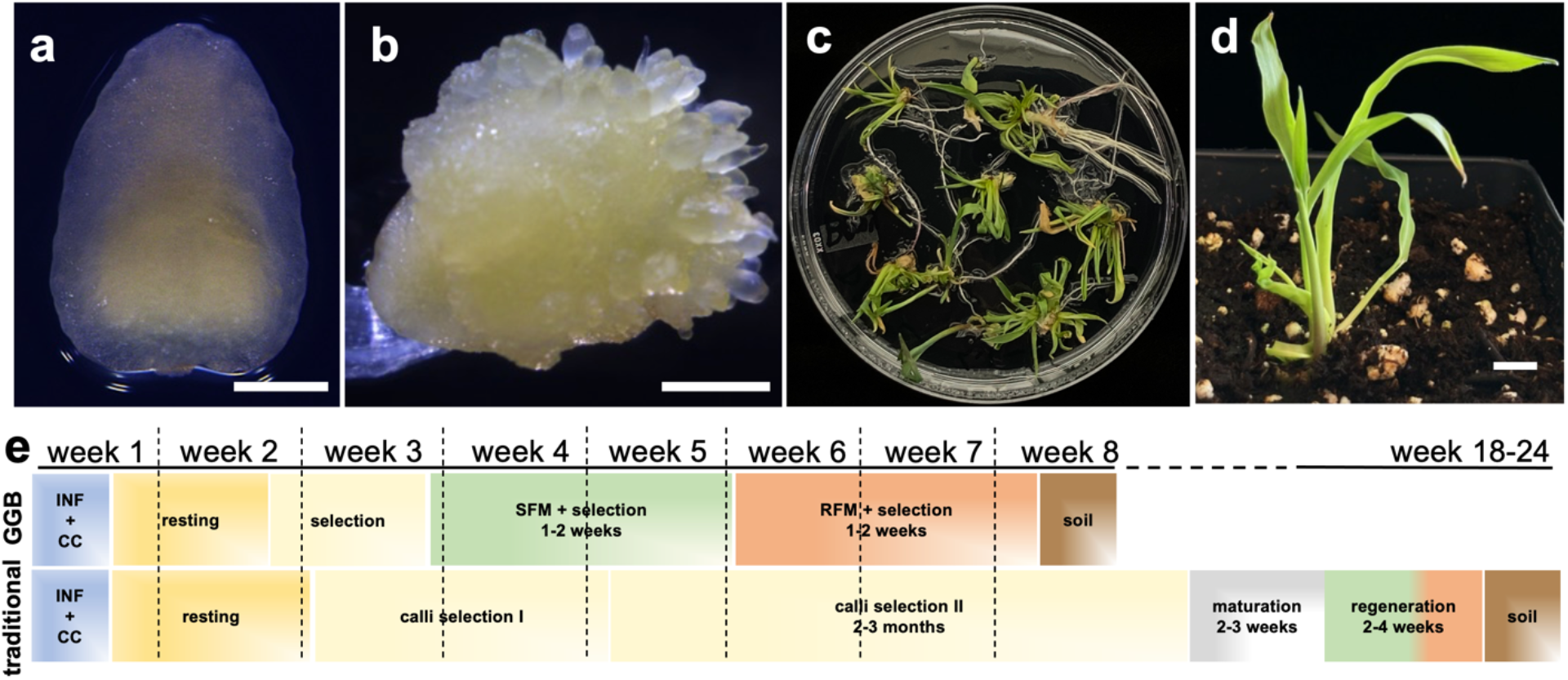
The GGB transformation system. (**a**) Immature embryo of the maize inbred line B104 used as explants. (**b**) Somatic embryos emerged on the surface of embryo scutellum side 6 days after *Agrobacterium* infection with the GGB construct. (**c**) Whole plantlet regeneration on rooting medium at 6 weeks after infection. (**d**) Regenerated plantlet was acclimated to soil in a growth chamber. (**e**) Timeline of the GGB transformation system compared to traditional transformation (INF + CC, infection and co-cultivation; SFM, shoot formation medium; RFM, root formation medium.

**Figure 3.**
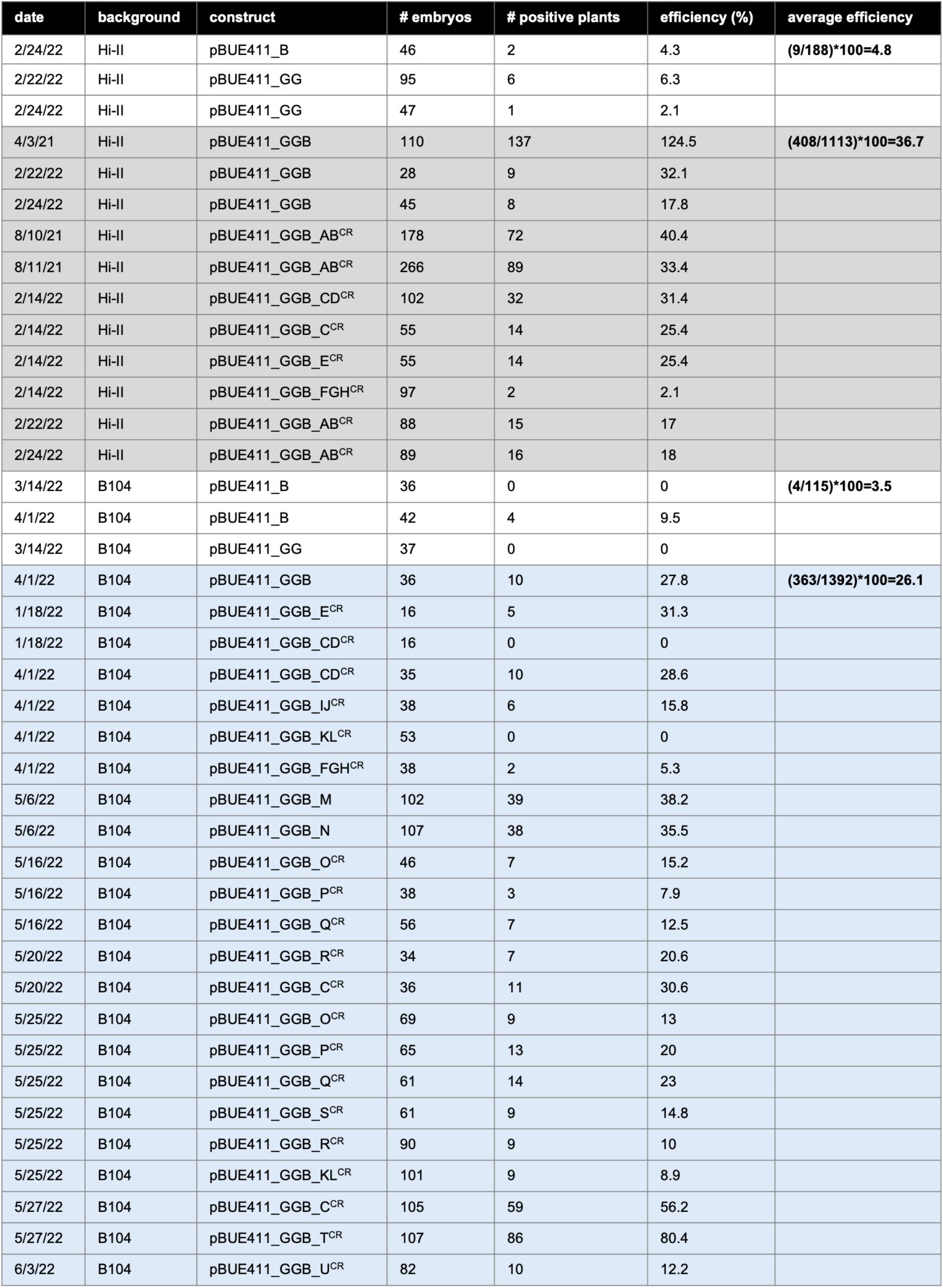
Efficiency of transformation in two different genetic backgrounds. Transformation efficiency is calculated as the number of treated embryos producing at least one confirmed transgenic plant, roughly corresponding to an independent transformation event (likely an underestimation since clusters of plants formed after regeneration are counted as individual plants). Letters (A-U) indicate targeted genes by CRISPR-Cas9, except M and N that indicate two fluorescent reporter constructs.

### The GGB system is highly efficient for maize embryo transformation

Several independent transformations were carried out on immature embryos of two commonly used maize transformation lines, Hi-II and B104 (Figure 3). Hi-II is a hybrid background (Mccaw *et al*. 2020), while B104 is an inbred background used by transformation facilities because of its vigor and better agronomic value relative to Hi-II (Raji *et al*. 2018; Kang *et al*. 2022). The complete list of experiments we conducted is provided in Figure 3. Transformation efficiency was calculated as the number of confirmed transgenic plants obtained after treating one embryo, roughly corresponding to an independent transformation event. We are likely underestimating efficiency since clusters of plants frequently formed that could not be easily separated after regeneration, and we counted the entire cluster as an individual plant. Regenerated plantlets were confirmed to be transgenic (carrying the pBUE411 vector with the *bar* resistant gene), first by applying the herbicide Liberty (0.2%) to a section of a mature leaf and scoring the resistance or sensitivity to the treatment after 5 days (also called BASTA painting technique), and with construct-specific PCRs (Figure 4). Overall, the efficiency of transformation was 37% and 26% in Hi-II and B104, respectively. Variability in all transformation experiments was still present (25.74±4.07; s.e.m.); however, only two attempts at transformation failed to produce transgenic plantlets, out of 34 total experiments. The efficiency of transformation in both genetic backgrounds was 7-fold higher than with using either component (BBM or GRF-GIF) in isolation (Figure 3; p=0.03), indicating that the synergistic interactions of these morphogenic factors yield substantial improvement to maize transformation.

**Figure 4.**
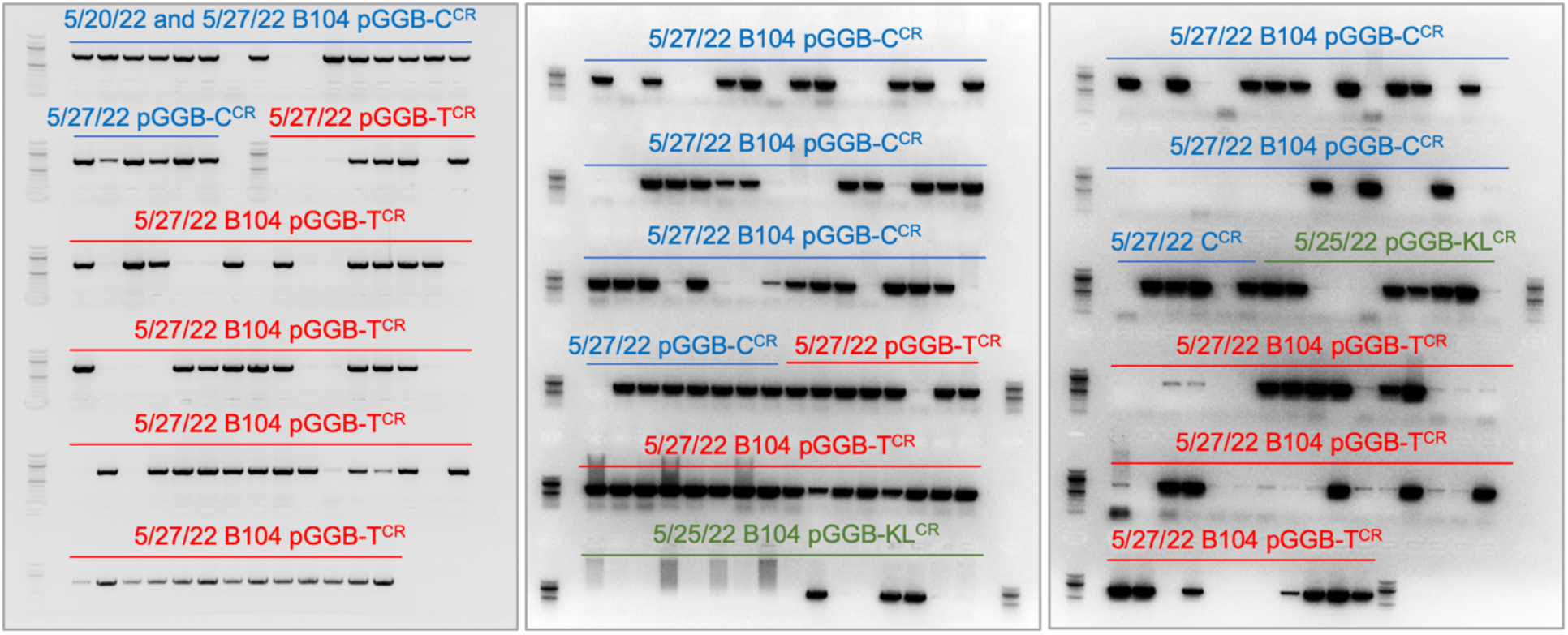
Genotyping of regenerated plantlets. After transplanting, plants were genotyped by PCR, using construct specific primers. An example of genotyping is shown for experiments of 5/20/22, 5/25/22 and 5/27/22.

### B104 plants containing the GGB vector do not show any obvious developmental defect

For monitoring potential effects on development by the two morphogenetic components of the GGB system, we focused on results from the B104 inbred line, a much vigorous and less variable background than Hi-II. No obvious effects on overall development and fertility were noted in the transformed plants carrying the GGB construct in both greenhouse and field conditions (Figure 5a-e). We compared plant height, leaf number and tassel branch number in T0 plants carrying the pBUE411-GGB construct and T0 plants without the construct (escapes from the regeneration protocol). No significant difference was observed for those traits, though T0 plants, independently of the presence of the construct, were in general slightly shorter and with fewer tassel branches than untransformed B104 plants (Figure 5f-h). However, these effects may be due to the competition for resources from plants growing in small clusters (see above).

**Figure 5.**
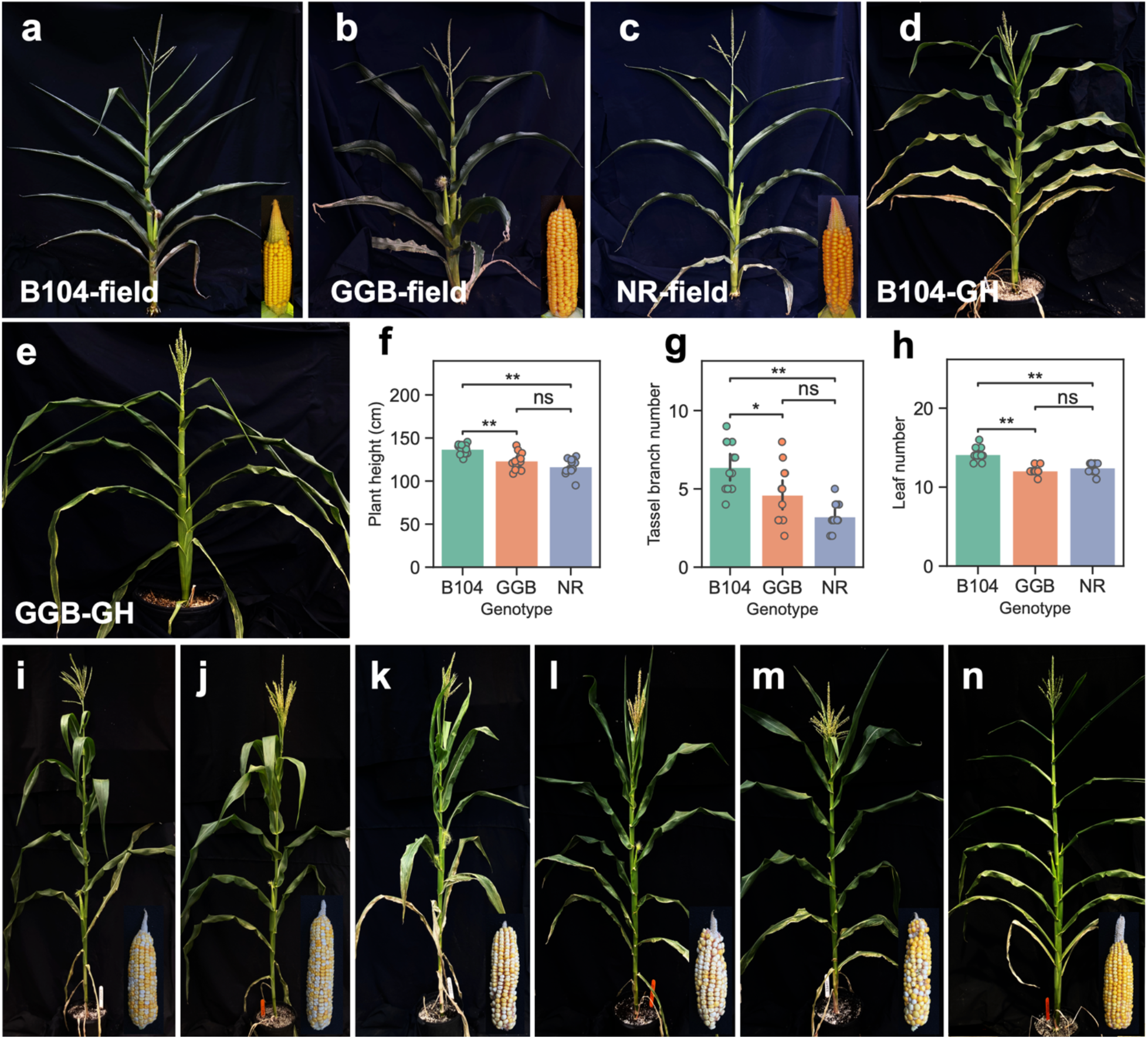
Mature plant phenotype of regenerated plants. (**a-e**) Vigorous B104 non-transformed (**a, d**) and regenerated plants (**b**,**c**,**e**) were grown in both greenhouse and field conditions. In (**c**) a non-transgenic plant regenerated using the same protocol (escape; NR, non-resistant to BASTA painting). (**f-h**) Quantification of T0 plants phenotype. t-test for independent samples with Bonferroni correction (n=13): (**f**) B104 vs. GBB p=6.706e-04, GGB vs. NR (non resistant) p=2.344e-01, B104 vs. NR p=1.745e-06; (**g**) B104 vs. GGB p=3.957e-02, GGB vs. NR p=5.737e-02, B104 vs NR p=3.479e-06; (**h**) B104 vs. GGB p=1.030e-07, GGB vs NR p=4.342e-01, B104 vs. NR, p=3.789e-05. ns, non significant; *, 1.00e-02 < p < 5.00e-02; **, p < 1.00e-02. (**i-n**) Mature T1 plants and ears resulting from crosses between T0 plants (Hi-II) and different inbred lines. Non-resistant (**i**; absence of *Bar* gene) and resistant (**j**; presence of *Bar* gene) plants of A619 X GGB (Hi-II), non-resistant (**k**) and resistant (**l**) plants of B73 X GGB (Hi-II), and non-resistant (**m**) and resistant (**n**) plants of B104 X GGB (Hi-II) crosses.

As for Hi-II transgenic plants carrying the GGB vector, given the variability in T0 plants that prevented a proper phenotypic evaluation, we verified that T1 plants originated from crosses between Hi-II T0 plants and different inbred backgrounds appeared as healthy and fertile as plants generated by the same cross but not carrying the construct (Figure 5i-n).

## DISCUSSION

The GGB system developed in this study represents a simple, rapid, and highly efficient method for maize transformation in both Hi-II hybrids and the inbred background B104. However, due to the inherent complications of the Hi-II background (mixed genetic background with lack of vigor), we recommend using the B104 inbred line for which we reached a respectable efficiency of transformation (26%). Even though variability in transformation efficiency is still observed among experiments and it is inherent to transformation experiments (see Figure 3, two experiments did not yield any transformants), nonetheless this system improves on the published efficiency of 6% obtained in B104 using the QuickCorn protocol and a ternary vector system (Kang *et al*. 2022). Given that we did not notice any detrimental effects on development, the GGB system could be suitable for large scale projects for the rapid generation of multiple transgenic lines, including fluorescent markers. Two of the experiments listed incorporated fluorescent reporters (pBUE411_GGB_M and pBUE411_GGB_N) with an efficiency above 30%, after removing the Cas9 cassette (details in Methods).

We believe that the transformation efficiency could be further improved by incorporating ternary vector systems (Anand *et al*. 2018; Zhang *et al*. 2019; De saeger *et al*. 2021; Kang *et al*. 2022; Wang *et al*. 2022b) that enhance virulence of *Agrobacterium* infections. We also anticipate that the GGB system with ternary vectors could improve transformation efficiency in different maize inbred lines, including recalcitrant lines such as B73, Mo17 and W22 (Springer *et al*. 2018; Sun *et al*. 2018; Hufford *et al*. 2021).

## METHODS

### Plant materials

The maize inbred B104 and Hi-II lines were grown in standard greenhouse conditions during winter and spring (2021-2022) and in summer nursery fields (2021) located at the Waksman Institute, in Piscataway, NJ.

### Maize GGB vector construction

We performed all PCR cloning with Phusion High-Fidelity DNA Polymerase (New England BioLabs). Total RNA was extracted from 3-5 mm ear primordia of the inbred line B73 using RNeasy Mini Kit (Qiagen) with on-column DNase I (Qiagen) treatment and used for complementary DNA (cDNA) synthesis with a ProtoScript® II First Strand cDNA Synthesis Kit (New England BioLabs). To clone the coding region of maize *BABY BOOM* (*BBM*, GRMZM2G141638) (Lowe *et al*. 2018), we performed PCR using cDNA generated from ear primordia, and the PCR product was cloned into the pENTR223 entry vector by SfiI digestion followed by sticky-end ligation performed by the T4 DNA ligase (New England BioLabs). To clone the 1098 bp promoter region and 5’-UTR of the PHOSPHOLIPID TRANSFERASE PROTEIN gene (*PLTP*, GRMZM2G101958) (Lowe *et al*. 2018), we performed PCR using genomic DNA extracted from leaves of B73, and the PCR product was cloned into pTF101 vector by HindIII and BamHI digestion followed by sticky-end ligation performed by the T4 DNA ligase. The coding region of maize *BBM* was then cloned into pTF101 vector containing the *PLTP* promoter sequence by LR Clonase II Enzyme Mix (ThermoFisher Scientific).

To generate the GGB construct containing BBM and the GRF4-GIF1 chimeric protein combined with Cas9 construct, we modified the available pBUE411 vector (Xing *et al*. 2014). In the first step, we performed PCR to clone *ZmUBIQUITIN*_pro_::*GRF4*-*GIF1* using primers JD633-F1/R1 and JD633 plasmid as templates(Debernardi *et al*. 2020). The resulting PCR product was cloned into pBUE411 vector at the PmeI cut site by Gibson Assembly using NEBuilder HiFi DNA Assembly Cloning Kit (New England BioLabs) and this produced the pBUE411_GRF-GIF construct (Figure 1a).

Subsequently, the guide RNA cassette of pBUE411 was replaced with a synthetic fragment carrying a multiple cloning site, and then the PLTP_pro_::BBM-NOS_term_ was amplified using the pTF101-PLTP_pro_::BBM-NOS_term_ plasmid as template and cloned into the multiple cloning site. The resulting pBUE411_GRF-GIF-BBM construct contains ZmUbi_pro_::GRF4-GIF1-NOS_term_, PLTP_pro_::BBM-NOS_term_ and ZmUbi_pro_::Cas9-E9_term_ (Figure 1a). To generate the pBUE411_BBM vector construct (pBUE411-PLTP_pro_::BBM-NOS_term_), the pBUE411_GRF-GIF-BBM construct was cut by AflII enzyme on double AflII cut sites to remove *ZmUBIQUITIN*_pro_::*GRF4*-*GIF1*, followed by self-ligation using T4 DNA ligase (New England BioLabs) (Figure 1a). The pBUE411_GGB_M and pBUE411_GGB_N constructs were created by removing the Cas9 cassette using AscI and NruI sites and the fluorescent reporter construct was inserted in those sites.

To incorporate the gRNAs into the pBUE411-GGB construct we first generated two entry vectors that carry OsU3_pro_-*Bsa*I_SmR_*Bsa*I-scaffold-OsU3_term_ and OsU6p2_pro_-*Bsa*I_SmR_*Bsa*I-scaffold-OsU3_term_, respectively (Figure 1b). Individual gRNA fragments were then ligated into the double *BsaI* sites by T4 DNA ligase following *BsaI* digestion to generate individual gRNA cassettes. Single or multiple individual gRNA cassettes were PCR amplified using primer combinations in the primer list (Table 1) and assembled into double *BsaI* sites of pBUE411-GGB construct (Figure 1a) using Golden Gate Assembly method. For Golden Gate Assembly reactions (15µl/reaction), equal molar concentrations of PCR products of gRNA cassettes and the pBUE411-GGB plasmid were mixed in a single 200µl Eppendorf tube. 0.5µl *BsaI* enzyme (New England BioLabs) and 1µl T4 DNA ligase (New England BioLabs) were then added to the tube, and finally, MilliQ water was added to bring the final volume to 15µl. Golden Gate Assembly reactions were setup in a PCR machine with 25-30 cycles of 37°C for 4 minutes and 16°C for 10 minutes. After the assembly reaction, the final plasmids were kept at 50°C for extra 5 minutes to linearize unexpected products carrying *BsaI* cut sites followed by 80°C for 10 minutes. 1 or 2µl of final reaction products were used for transformation of chemically competent *E. coli* cells.

**Table 1.**
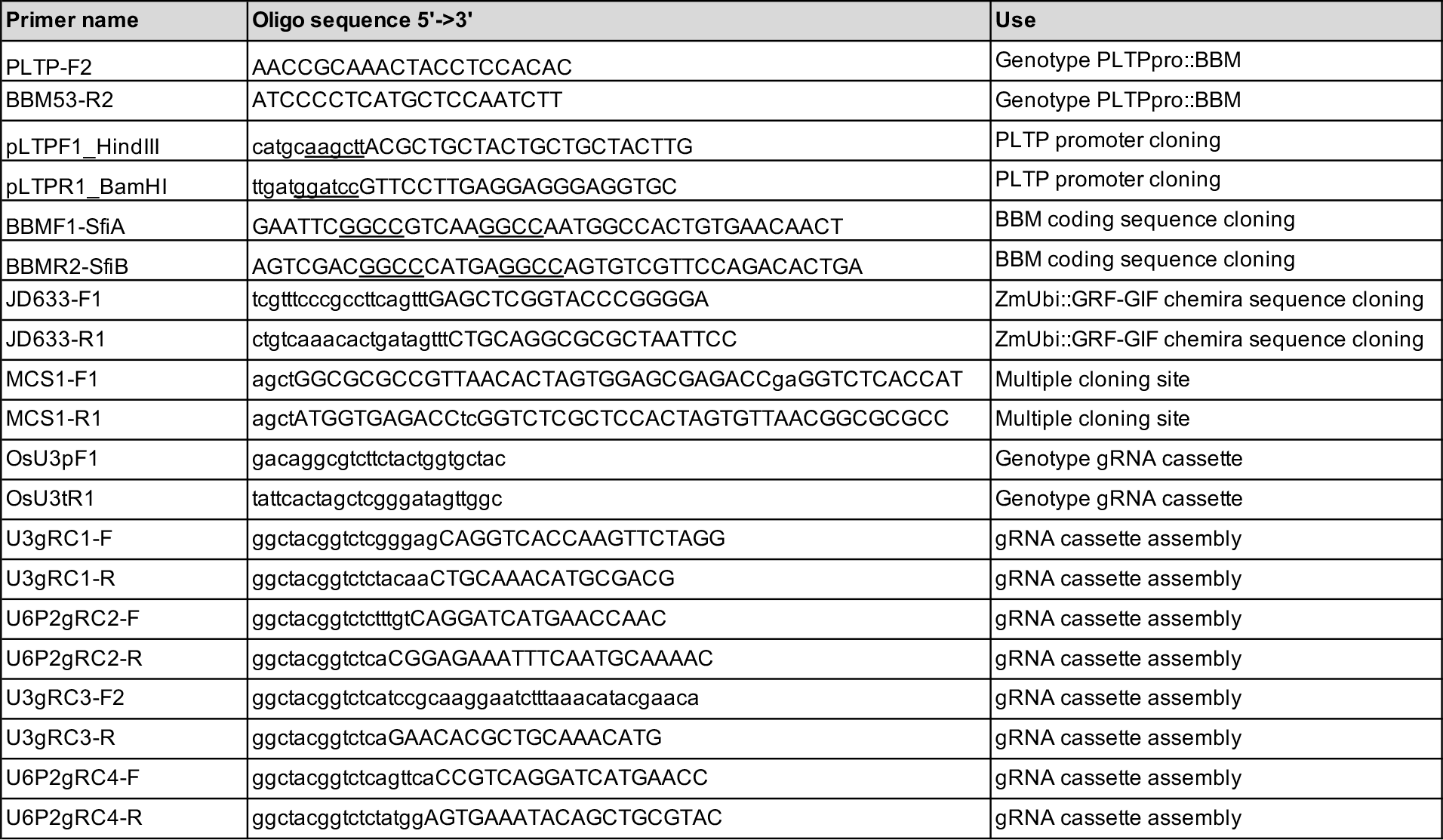
Primers used in this study.

### Maize transformation protocol

Maize transformation of immature embryos followed previously published protocols (Lowe *et al*. 2018; Masters *et al*. 2020) with a few modifications. For harvesting immature embryos, we grew Hi-II and B104 plants in standard greenhouse or in field conditions. We harvested fresh ears (10-14 days after pollination) and removed all husk leaves and silks, and then surface sterilized them for 20 minutes in 50% commercial bleach (1.65% sodium hypochlorite) with 0.1% of Tween-20. After surface sterilization, we washed the ears three times with sterilized water and dissected the immature embryos in laminar flow cabinets (embryo sizes, 1.5–2.0 mm) using a flame sterilized spatula. The fresh embryos were collected in a 2 ml microcentrifuge tube filled with infection medium (**700A**) and subsequently washed for three times with the same infection medium. For *Agrobacterium tumefaciens* inoculation, the *Agrobacterium* strain EHA105 stored as a glycerol stock at −70°C carrying the pBUE411 vector was grown on a **YP** agar plate containing 100 mg/liter kanamycin, 30 mg/liter rifampicin in the dark for 2 days at 28 °C. Two *Agrobacterium* colonies were picked and cultured in 15 ml falcon tubes filled with 5 ml YP liquid medium containing 100 mg/liter kanamycin, 30 mg/liter rifampicin and 200 µM acetosyringone overnight at 28 °C. The next day, 2 ml of *Agrobacterium* cells were pelleted in a centrifuge (8000g) for 3 minutes, and resuspended in the **700A** medium. The suspension culture was then transferred to 15ml tubes (4-6ml/tube) and adjusted to 0.35 – 0.4 at OD_600_ using a spectrophotometer. We gently shook this suspension culture in the dark for more than 2 to 6 hours at room temperature (21-25°C), and then immersed all the embryos in 1.8ml of *Agrobacterium* suspension with **700A** infection medium for 5 minutes. Infected embryos together with *Agrobacterium* suspensions were poured into **562V** co-cultivation plates, and the embryos were spread over the surface by gently shaking the plates by hand. The excess of *Agrobacterium* suspension was then removed from the plates by pipetting and the embryos with scutellum side up were incubated at 20 °C in the dark. After 3 days of co-cultivation, we transferred all embryos (maintaining the scutellum side up) onto the resting medium **605T** without selection and incubated them at 28 °C in the dark. After 7 days, we transferred all embryos to the **605T** selection medium with 5 mg/l bialaphos and incubated them at 28 °C in the dark. After an additional 7 days, we transferred embryos with somatic embryos on the scutellum side to the shoot formation medium (**13329A**) containing 5 mg/liter bialaphos. After an additional one to two weeks, we transferred the regenerated shoots with emerging leaves to rooting medium (**13158**) for 1-2 weeks at 28 °C in a light chamber (16h day/8h night, 20-150 µmol/m2/s) for 2 weeks. Rooted plants were acclimated to soil by transferring them to a tray filled with potting media, covering them with a clear plastic dome and maintaining them for 1 to 2 weeks in a Conviron GR64 growth chamber (16h day/8h night, 20-150 µmol/m2/s) at 28 °C. As the plants became more vigorous, we then transplanted individual plants into 5-gallon (18.9 l) pots containing pre-wetted soil and maintained them in the greenhouse in standard growing conditions. We verified positive transformed plants by using light applications of Liberty herbicide (0.3%) on 3^rd,^-5^th,^ leaves (“BASTA painting”), and scoring resistance (presence or absence of the *Bar* gene) after approximately 5 days and by construct specific PCRs (Table 1).

## RECIPES

### YP Agar

Peptone, 10 g/l

Yeast Extract, 5 g/l

NaCl, 5 g/l

pH to 6.8 with NaOH

Add Bactoagar, 13 g/l

Autoclaved for 15 minutes, cool to 55^0,^C and added:

Kanamycin, 50 mg/l

Rifampicin, 30 mg/l

### Infection medium: 700A

MS Basal Medium, 4.4 g/l (PhytoTech Labs)

2,4-D, 1.5 mg/l (PhytoTech Labs)

Sucrose, 68.5 g/l

Glucose, 36 g/l

pH to 5.8 with NaOH

Autoclaved for 15 minutes.

### Co-cultivation medium: 562V

N6 Basal Salt Mixture, 4.0 g/l (PhytoTech Labs)

2,4-D, 2.0 mg/l (PhytoTech Labs)

Sucrose, 30 g/l

pH to 5.8 with NaOH, then add Agar, 8 g/L

Autoclaved, cooled to 55°C and added:

Silver Nitrate, 1 mg/l

Acetosyringone, 100 µM

Poured into 15×100 Petri dishes, 30 ml/plate and let them solidify for 30 minutes and stored at 4 °C.

### Resting medium: 605T

MS Basal Salt Mixture (1x), 4.3 g/l (PhytoTech Labs)

N6 Macro Salts (0.6x), 60 ml/l

B5 Micro Salts (0.6x), 0.6 ml/l

Eriksson’s Vitamins (0.4x), 0.4 ml/l (PhytoTech Labs)

S&H Vitamins (0.6x), 0.6 g/l (PhytoTech Labs)

Ferrous Sodium stock (0.6x), 6 ml/l

KNO_3_, 1.68 g/l

Thiamine HCl, 0.2 mg/l

Casein Hydrolysate, 0.3 g/l (PhytoTech Labs)

2,4-D, 0.8 mg/l (PhytoTech Labs)

L-proline, 2 g/l (PhytoTech Labs)

Sucrose, 20 g/l

Glucose, 0.6 g/l

pH to 5.8 with NaOH, then added phytagel, 2.5 g/l

Autoclaved, cooled to 55 ^o,^C and added:

Dicamba, 1.2 mg/l

Cefotaxime, 100 mg/l

Timentin, 100 mg/l

Silver Nitrate, 3.4 mg/l

Poured into 15×100 Petri dishes, 30 ml/plate and let them solidify for 30 minutes and stored at 4 °C.

### Shoot Formation: 13329A

MS Basal Medium, 4.4 g/l (PhytoTech Labs)

Zeatin, 0.5 mg/l (PhytoTech Labs)

Sucrose, 60 g/l

pH to 5.8 with NaOH, then added phytagel, 2.5 g/l

Autoclaved, cooled to 55^o,^ C and added:

Thidiazuron, 0.1 mg/l (PhytoTech Labs)

BAP, 1 mg/l (PhytoTech Labs)

Carbenicillin, 100 mg/l

5 mg/l Bialaphos

Poured into 15×100 Petri dishes, 30 ml/plate and let them solidify for 30 minutes and stored at 4 °C.

### Rooting medium: 13158

MS Basal Medium, 4.4 g/l (PhytoTech Labs)

Sucrose, 40 g/l

pH to 5.8 with NaOH, then added phytagel, 2.5 g/l

Autoclaved, cooled to 55 ^o,^C and added:

Carbenicillin, 100 mg/l

5 mg/l Bialaphos

Poured into 25×100 Petri dishes, 30 ml/plate and let them solidify for 30 minutes and stored at 4 °C

## DATA AVAILABILITY

The GGB vector is currently available upon request.

## ACKNOWLEDGMENTS

We acknowledge funding from the National Science Foundation (IOS #1546873 and 1916804) to A.G.

## AUTHOR CONTRIBUTIONS

Z.C. and A.G. designed research and analyzed data; Z.C. performed experiments; J.M.D. and J.D. provided material for vector construction; Z.C. and A.G. wrote the paper with contributions from all authors; all authors approved the manuscript.

